# An enriched network motif family regulates multistep cell fate transitions with restricted reversibility

**DOI:** 10.1101/453522

**Authors:** Yujie Ye, Jordan Bailey, Chunhe Li, Tian Hong

**Affiliations:** Department of Biochemistry and Cellular and Molecular Biology, The University of Tennessee,Knoxville, Tennessee, United States of America; Shanghai Center for Mathematical Sciences, Fudan University, Shanghai, China; Institute of Science and Technology for Brain-Inspired Inteligence, Fudan University, Shanghai; National Institute for Mathematical and Biological Synthesis, Knoxville, Tennessee, United States of America

## Abstract

Multistep cell fate transitions with stepwise changes of transcriptional profiles are common to many developmental, regenerative and pathological processes. The multiple intermediate cell lineage states can serve as differentiation checkpoints or branching points for channeling cells to more than one lineages. However, mechanisms underlying these transitions remain elusive. Here, we explored gene regulatory circuits that can generate multiple intermediate cellular states with stepwise modulations of transcription factors. With unbiased searching in the network topology space, we found a motif family containing a large set of networks can give rise to four attractors with the stepwise regulations of transcription factors, which limit the reversibility of three consecutive steps of the lineage transition. We found that there is an enrichment of these motifs in a transcriptional network controlling the early T cell development, and a mathematical model based on this network recapitulates multistep transitions in the early T cell lineage commitment. By calculating the energy landscape and minimum action paths for the T cell model, we quantified the stochastic dynamics of the critical factors in response to the differentiation signal with fluctuations. These results are in good agreement with experimental observations and they suggest the stable characteristics of the intermediate states in the T cell differentiation. These dynamical features may help to direct the cells to correct lineages during development. Our findings provide general design principles for multistep cell linage transitions and new insights into the early T cell development. The network motifs containing a large family of topologies can be useful for analyzing diverse biological systems with multistep transitions.

**Author summary:** The functions of cells are dynamically controlled in many biological processes including development, regeneration and disease progression. Cell fate transition, or the switch of cellular functions, often involves multiple steps. The intermediate stages of the transition provide the biological systems with the opportunities to regulate the transitions in a precise manner. These transitions are controlled by key regulatory genes of which the expression shows stepwise patterns, but how the interactions of these genes can determine the multistep processes were unclear. Here, we present a comprehensive analysis on the design principles of gene circuits that govern multistep cell fate transition. We found a large network family with common structural features that can generate systems with the ability to control three consecutive steps of the transition. We found that this type of networks is enriched in a gene circuit controlling the development of T lymphocyte, a crucial type of immune cells. We performed mathematical modeling using this gene circuit and we recapitulated the stepwise and irreversible loss of stem cell properties of the developing T lymphocytes. Our findings can be useful to analyze a wide range of gene regulatory networks controlling multistep cell fate transitions.

## Introduction

Cell fate transition, including differentiation, de-differentiation and trans-differentiation, is a fundamental biological process in which the function of a cell gets specialized, reprogrammed or altered. The process often involves significant changes of multiple cellular properties, including the morphology, the self-renewal capacity and the potentials to commit to alternative lineages [1,2]. These changes are controlled by the dynamics of interacting transcription factors (TFs) and the modulation of chromatin structures, which in term are governed by complex regulatory networks in the cells [3-5]. Interestingly, the fate transitions in many systems are achieved by sequential commitments to a series of cellular states with stepwise changes in their transcriptional profile towards the final stage of the program (Figure 1) [6-11]. The intermediate states between the initial state (e.g. the undifferentiated state in the case of cell differentiation) and the final state may be important for multiple purposes, such as facilitating ‘scheckpoints’ that ensure appropriate development of cellular behaviors, or allowing the cells to make correct decisions at the lineage branching points [11-15].

**Figure 1.**
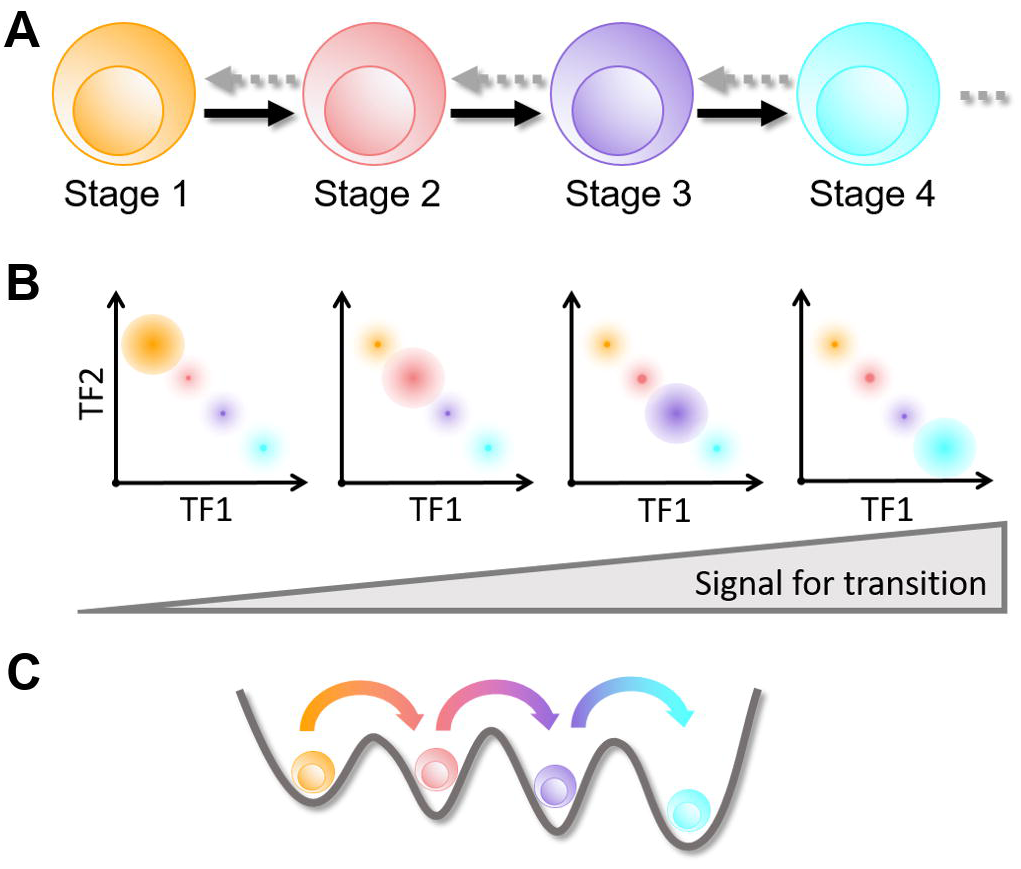
Illustration of multistep cell fate transition. **A.** transition from one cellular state to another via two intermediate states. Dashed arrow indicates the limited reversibility of each transition. **B.** stepwise changes of the levels of two transcription factors during the multistep transitions involving four states. **C.** metaphoric energy landscape depicting the four-attractor system. Colors for cell states and transition arrows in B and C match those in the illustration in A.

One example of these stepwise cell lineage transitions is the development of T lymphocytes in the thymus. The differentiation from multipotent pre-thymic progenitor cells to committed T cells involves multiple cellular states with stepwise changes of their cellular properties and the transcriptional profiles (Table 1) [16-19]. Several lines of evidence suggest that the transition states at an early phase of the differentiation can serve as stable checkpoints for sequential lineage commitments. The progress through these intermediate states is accompanied by stepwise loss of their potentials to differentiate into other cell types: pre-thymic progenitor cells can be converted to a few types of cells, including B cells, natural killer (NK) cells, dendritic cells (DCs) etc., whereas the multipotency of the intermediate cell types is more limited but not completely lost [20-26]. In addition, the stability of these intermediate states is substantial because the loss of differentiation signals does not result in de-differentiation of some intermediate states [20], suggesting restricted reversibility (or complete irreversibility) of the multiple transitions. In addition, the lymphoid progenitor cells need to divide for a certain number of times at an intermediate state before committing to the T cell lineage, and the stable activities of the lineage defining transcriptional program at the intermediate stages may be important for the proliferations [27]. Finally, the loss of certain transcription factors (e.g. BCL11B) can lead to the termination of the differentiation at some intermediate states, which is often associated with diseases such as leukemia [18,20,28]. This further suggests that the intermediate states are cellular ‘attractors’ between the initial and the final stages of the differentiation (Figure 1, bottom panel). Similar stable intermediate states during cell lineage transitions are observed in other systems, such as the epithelial-mesenchymal transition, and the skin development (Table 1), and those states also serve as regulatory stages for altering cellular properties including self-renewal and migration [10,29-37]. Therefore, the multiple intermediate states are involved in diverse normal development and pathological conditions. Understanding the regulatory programs for the sequential cell lineage commitments is a key step towards the elucidation of mechanisms underlying various biological processes involving multistep lineage transitions. Despite the accumulating data and observations on these stepwise lineage commitments, general mechanisms governing these differentiation processes with multiple intermediate cellular states remain unclear.

**Table 1.** Examples of multistep transitions with restricted reversibility

| Physiological scenario | Cellular phenotypic transition | Key regulators with stepwise modulations | Extracellular signals | Evidence supporting multistep transitions, multiple intermediate states and restricted reversibility |
| --- | --- | --- | --- | --- |
| Early T cell development | ETP/DN1 → DN2a → DN2b → DN3 | PU.1 TCF-1<br>GATA3<br>BCL11B<sup>b</sup> | Notch | [17-20,28] |
| Skin development | Stem cell → renewable spinous cell → non-renewable spinous cell → granular cell | OVOL1 OVOL2 | Calcium ion | [32,33,37] |
| Epithelial-mesenchymal transition | E →<sup>a</sup> EM1 →<sup>a</sup> EM2 →<sup>a</sup> M | SNAIL1 TWIST<br>ZEB1 miR200 | TGF-β | [29-31,34-36] |

<sup>a</sup> Reversal transitions were observed, but they occur in a limited subpopulation.98 <sup>b</sup> Unlike other factors, BCL11B exhibits an abrupt change at the second transition.

In this study, we explored the strategies in terms of the transcriptional network design that gives rise to stepwise transitions during cell differentiation. We first used a generic form of networks containing three interacting TFs to find network motifs that can produce four attractors (the minimum number of attractors in the examples of T cell development, epithelial-mesenchymal transition and skin development) with stepwise changes of transcriptional factor levels. We found two types of network motifs, both involving interconnections of positive feedback loops, which can generate the four-attractor systems. These motifs constitute a large family of gene regulatory networks. We found that there is an enrichment of these motifs in a network controlling the early T cell development. We built a specific model using known interactions among key transcription factors in developing T cells, and the model shows that the transcriptional network governs multistep and irreversible transitions in the development process.

To investigate the stochastic dynamics for early T cell development model, we mapped out the quasi-energy landscape for the early T cell development. This landscape characterizes the four attractors representing four stages of early T cell development quantitatively. In addition, by calculating the minimum action paths (MAPs) between different attractors, we quantified the dynamics of the key factors in response to Notch signal with fluctuations, which are in good agreement with experimental observations. Finally, we identified the critical factors influencing T cell development by global sensitivity analysis based on the landscape topography. Overall, our model for early T cell development elucidates the mechanisms underlying the stepwise loss of multipotency and multiple stable checkpoints at various stages of differentiation. The network topologies for multiple attractors found in this study and our motif discovery strategy combined with the landscape methodology can be useful for analyzing a wide range of cell differentiation systems with multiple intermediate states.

## Results

### Networks in a large motif family govern systems with four attractors with stepwise transcriptional modulation

To find transcriptional network topologies that can generate multiple intermediate states during cell fate transition, we first performed random parameter sampling with a network family containing up to 3 nodes (Figure 2A). In this framework of network topology, each node represents a transcription factor (TF) that can potentially influence the transcription levels of other two TFs and itself. Topology searching with a 3-node network was used for motif discovery for various performance objectives in previous studies [38,39]. We performed exhaustive search for topologies with up to 6 regulations from a total of 9 regulations of the network family, and constructed a mathematical model for each topology (see Methods for details). For each model, we performed random sampling of the parameter space from uniformly distributed values (Table S1). and we selected topologies containing at least one parameter set that is able to generate four attractors with stepwise changes of transcriptional levels. We define the system with four attractors with the stepwise changes of transcriptional levels as the scenario in which there are four stable steady states and they can be consistently ordered by the concentrations of any pairs of TFs. In other words, one TF always monotonically increases or decreases with another TF in these four states. Among the 2114 network topologies that we searched, we found 216 topologies that can produce such behavior. In addition, we found 417 topologies that can only produce four unordered steady states (TF concentrations are non-monotonically correlated among the states) (Figure S11).

**Figure 2.**
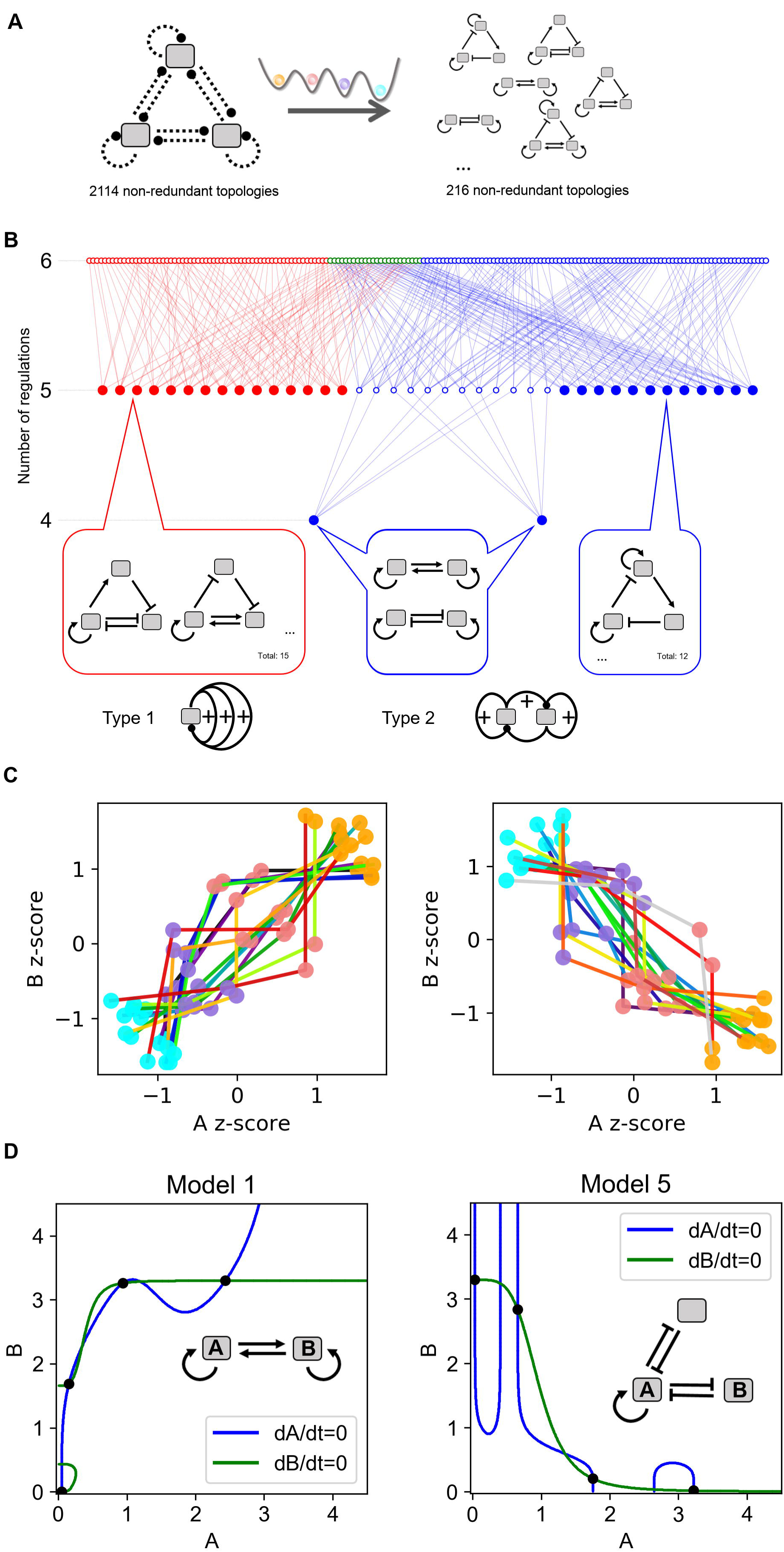
Network motifs governing four-attractor systems. **A.** Illustration of the network topology searching. Dashed arrows are regulations sampled. The topologies were screened by the criterion of the four attractors with stepwise changes of TFs. **B.** Complexity atlas for selected topologies. Closed circles denote minimum motifs. Open circles denote topologies containing more regulations than those in the minimum motifs. Each arrow denotes the difference by one regulation in the network. Examples of minimum motifs are shown at the bottom. Red: Type I motif. Blue: Type II motif. Green: Hybrid motif. **C.** Overlaid four attractors for each of the 29 minimum topologies. Factor A denotes the TF on the left of the network diagram. Factor B denotes the TF on the right of the network diagram. In some topologies A and B and positively correlated (left panel), whereas they are negatively correlated in other topologies (right panel). Colored dots denote the stable steady states. Colored lines connect states of their corresponding topologies. The colors of the cell states match the illustration in Figure 1. The colors of the lines denote different representative models. z-score is calculated by shifting the mean of each four attractors to 0 and then normalizing the four data points to unit variance data. **D.** Example phase planes for two minimum topologies (Type I and Type II respectively). In each case, four out of the seven steady states (intersections denoted by solid dots) are stable. Network structures and phase planes for all 29 minimum motifs are included in Figures S1 and S2. All models shown in this figure are built with additive form of Hill functions.

To visualize the relationships among these topologies, we constructed a complexity atlas (Figure 2B), in which the nodes represent the network structures that gave rise to four attractors, and the edges connect pairs of topologies that differ by a single regulation (addition or removal of a transcriptional interaction) [40]. We define the minimum topologies as those of which the reduction of complexity, or the removal of any regulation from the network, will abolish its capability to generate four attractors (solid nodes in Figure 2B and examples in Figure 2C). We found 29 such minimum topologies which represent the non-redundant structures for producing the four-attractor system.

Interestingly, all of the 216 topologies obtained from our search contain three distinct positive feedback loops (including double-negative feedback loops), and they can be categorized into two types of motifs (Figure 2B, bottom panel). The Type I motif contains three positive feedback loops that are closed at a single TF (red nodes and edges in Figure 2B). The Type II motif contains three connected positive feedback loops, two of which do not share any TF but are connected via the third loop (blue nodes and edges in Figure 2B). There is a remarkable diversity of each of the motif types because the interconnected positive feedback loops can share multiple TFs (Figures S1 and S2). Based on the complexity atlas (Figure 2B), we found that Type II motifs contain 4-6 regulations, and Type I motifs contain 5-6 regulations. Some of the networks with 6 regulations contain subnetworks of both Type I and Type II motifs (Hybrid type, green nodes). The four attractors in the space of two TFs exhibit a variety of patterns of nonlinear monotonic correlations (Figure 2C, Figure S3), which are governed by intersections of highly nonlinear nullclines in the state space containing the two TFs (Figure 2D, Figures S1 and S2). The definitions of various types of motifs are listed in Table 2, and the statistics of the topologies discovered are summarized in Table 3 (also see Figure S11 for an illustration). Overall, this motif family represents a large number of networks that can produce a common type of behaviors: multiple stable intermediate states in terms the transcriptional activity.

**Table 2.** **Definitions and key features of network motifs that generate systems with four ordered attractors.**

|  | Definition | Minimum number of regulations | Minimum number of positive feedback loops |
| --- | --- | --- | --- |
| Type I motifs | Three positive feedback loops that share one or more TFs among all of them. | 5 | 3 |
| Type II motifs | Three connected positive feedback loops. Two of them do not share any TF but are connected via the third loop. | 4 | 3 |
| Hybrid motifs | Motifs containing both Type I and Type II motifs. | 6 | 4 |

**Table 3.**
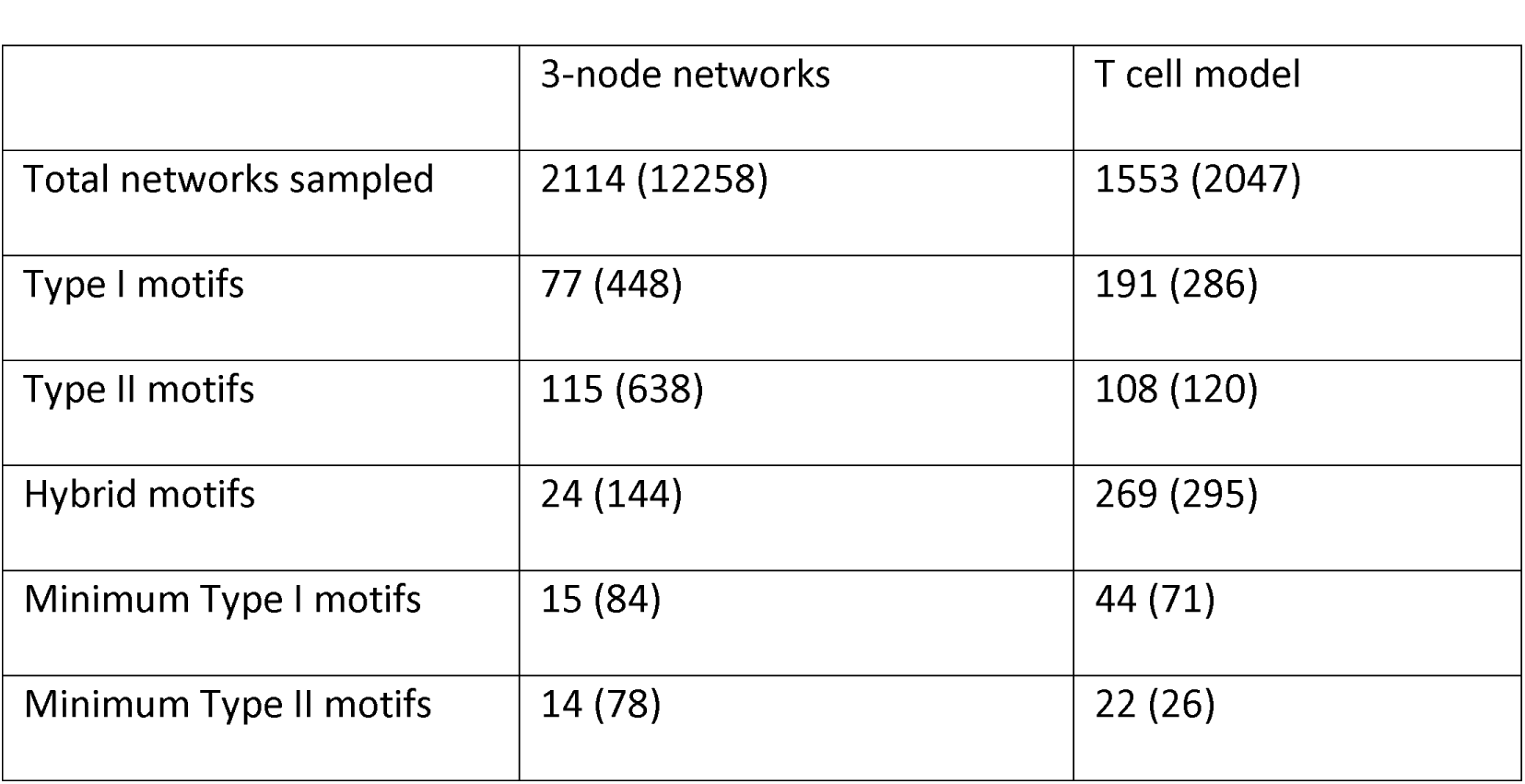
**Numbers of sampled network structures and discovered motifs ^a^**

|  | 3-node networks | T cell model |
| --- | --- | --- |
| Total networks sampled | 2114 (12258) | 1553 (2047) |
| Type I motifs | 77 (448) | 191 (286) |
| Type II motifs | 115 (638) | 108 (120) |
| Hybrid motifs | 24 (144) | 269 (295) |
| Minimum Type I motifs | 15 (84) | 44 (71) |
| Minimum Type II motifs | 14 (78) | 22 (26) |

<sup>a</sup>In each cell of the table, the first number is the number of non-redundant network topologies. The number in the parentheses is the number of networks (or sub-networks of the T cell model) including the isometric topologies.

In summary, we found two types of network motifs that generate four attractors with stepwise changes of the transcriptional profile. Two of these attractors represent the multiple intermediate states observed in various biological systems. This exploratory analysis elicits several interesting questions: what are the biological examples of such network motifs? Can the conclusions with respect to the two types of motifs be generalized to networks with more than three TFs? Is there any advantage of combining both types of motifs? How are the transitions among these states triggered deterministically and stochastically? To provide insights into these questions in a more biologically meaningful context, we will use a specific biological system to describe more detailed analysis of these motifs and their underlying gene regulatory networks in the following sections.

### Type I and Type II network motifs are enriched in a transcriptional network controlling early T cell development

We asked whether the motifs that we discovered can be found in any known transcriptional network that potentially control multistep cell differentiation. We used the early T cell differentiation in the thymus as an example to address this question. The differentiation from multipotent lymphoid progenitor cells to unipotent early T cells involves multiple stages at which the cells possess varying potentials to commit to non-T lineages and other cellular properties such as proliferation rates. At the early phase of this process, four stages of development T cells (ETP/DN1, DN2a, DN2b, DN3) were identified experimentally, and the progression through these stages is controlled by a myriad of transcription factors including four core factors, TCF-1, PU.1, GATA3 and BCL11B. These TFs form a complex network among themselves (see Figure 3A and supporting experimental observations in Table S3), and the stepwise changes in the levels of these TFs were observed in the four developmental stages of T cells [20,28]. The interactions involving these core TFs were shown to be critical for the irreversible commitment to the T cell lineage by forming a bistable switch [41]. Among these factors, PU.1 level decreases as the cells commit to later stages, whereas the levels of other three factors increase in this process. It is unclear, however, whether this transcriptional network can serve as a regulatory unit that governs the multistep nature of the T cell differentiation.

**Figure 3.**
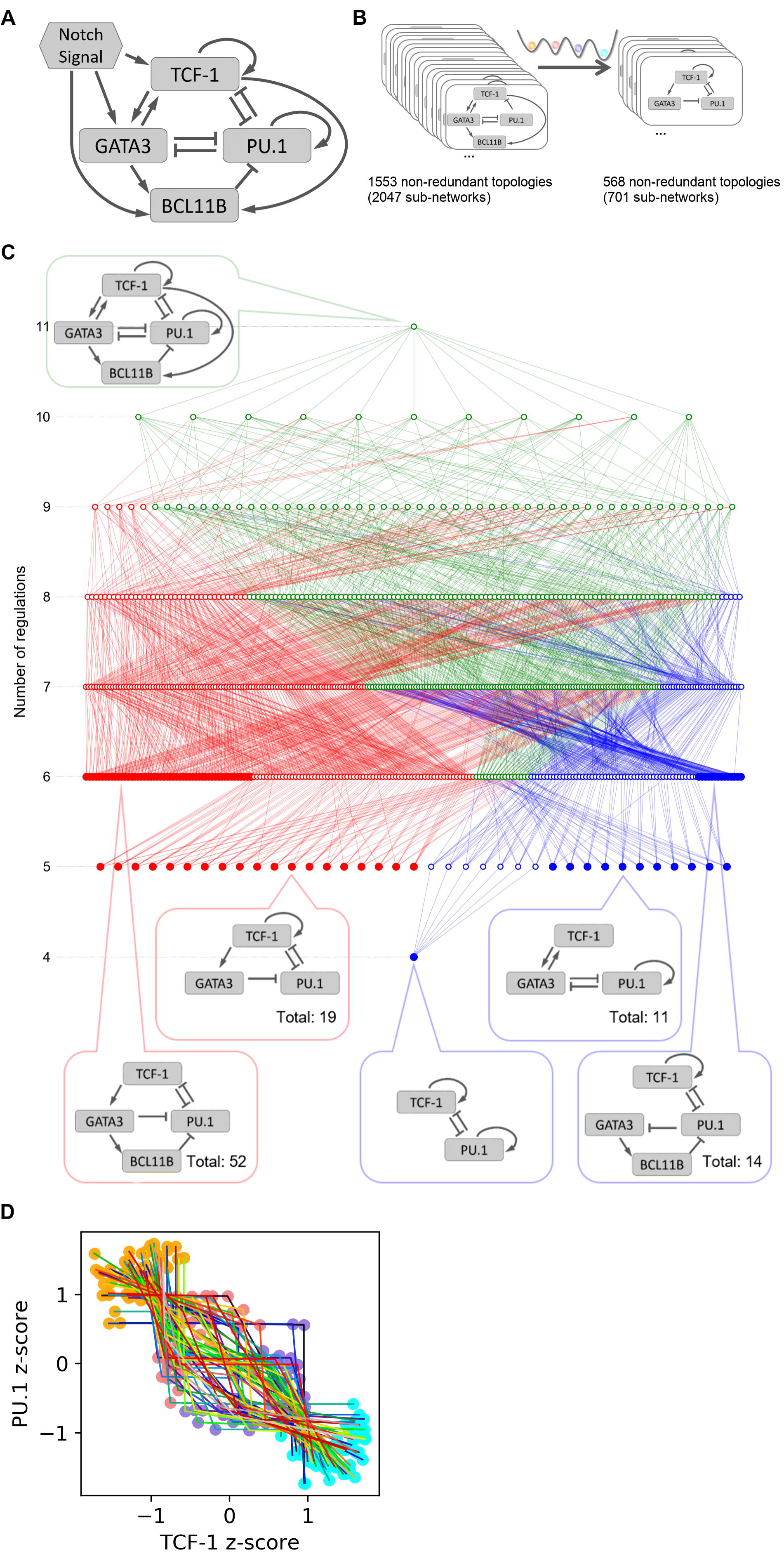
Four-attractor motifs in the early T cell transcriptional network. **A.** Influence diagram for transcriptional regulations among four core factors controlling the early T cell development. **B.** Functional subnetworks of the T cell network were systematically obtained by removing regulations from the network. These subnetworks were screened by the criterion that four attractors with stepwise changes of TFs exist in the absence of Notch signal. **C.** Complexity atlas showing the relationships of the two four-attractor motifs in the subnetworks of the T cell model. Top callout shows the full network in the absence of Notch. Bottom callouts show examples of the minimum functional subnetworks of the two types with particular numbers of regulations. Red: Type I motif. Blue: Type II motif. Green: Hybrid motif. **D.** Overlaid four attractors for each of the 66 minimum topologies. Colored dots denote the stable steady states. Colored lines connect states of their corresponding topologies. All models shown in this figure are built with the multiplicative form of Hill functions.

We noticed that this T cell transcriptional network contains the motifs that we found in our analysis using the generic form of networks, we therefore hypothesized that the models based on this network can have four attractors with sequential changes of the four TFs. Indeed, using random sampling we were able to find parameter sets that give rise to four-attractor systems similar to what we obtained with the generic 3-node framework. To find the functional components that generate this behavior, we analyzed the subnetworks of the complex T cell regulatory network [42]. We removed the regulations from the network systematically, and we found that out of the non-redundant 1553 topologies (2047 subnetworks), there are 568 topologies (701 subnetworks) that can generate four attractors with stepwise changes of the TFs (Figure 3B). We used a complexity atlas to visualize the relationships among these subnetworks (Figure 3C). We found that the network can be reduced to one of the 66 minimum topologies (97 minimum subnetworks) which retains the four-attractor property (solid nodes in Figure 3C). Notably, these networks can be classified into the two types of motifs described earlier (Figure 2B). Similar to the networks that we obtained through the generic framework, the two types of minimum motif have 4-6 regulations. Subnetworks with both types of motifs (green nodes and edges) start to appear when the number of regulations reaches six. The numbers of motifs and subnetworks obtained for the generic framework and the T cell model are summarized in Table 3.

We next quantified the enrichment of the two motif families in the early T cell transcription network. We first generated random networks by perturbing the existing regulations in the network model and computed the empirical p-values for observing the numbers of different types of network motifs. The T cell network contains a large number of positive feedback loops and the two types of motifs that we described earlier (Figure 4, top panel). As expected, the network is significantly enriched with positive feedback loops in general (Figure 4, middle panel, red bars). However, the enrichments of Type I motifs and the combinations of Type I and Type II motifs are even more significant than that of the single positive feedback loops (Figure 4, middle panel, red bars). To exclude the possibility that this differential significance was observed due to the way we generate random networks which gives low p-values (<10^−4^) in general, we used another method to generate random networks with an augmented number of regulations (Figure 4, middle panel, blue bars). Each pair of TFs were assigned with a pair of random regulations (positive, negative or none). Consistent with the previous method, the T cell transcriptional network is enriched with positive feedback loops overall, but the enrichment is more significant for Type I motifs or for the combination of Type I and Type II motifs. Interestingly, motifs that are similar to Type I motif but have higher complexity (more positive feedback loops) does not show more significant enrichment than Type I motif does (Figure S12). These results suggest the possibility that the network has been evolved to reach more complex performance objectives than those enabled by simple positive feedback loops alone.

**Figure 4.**
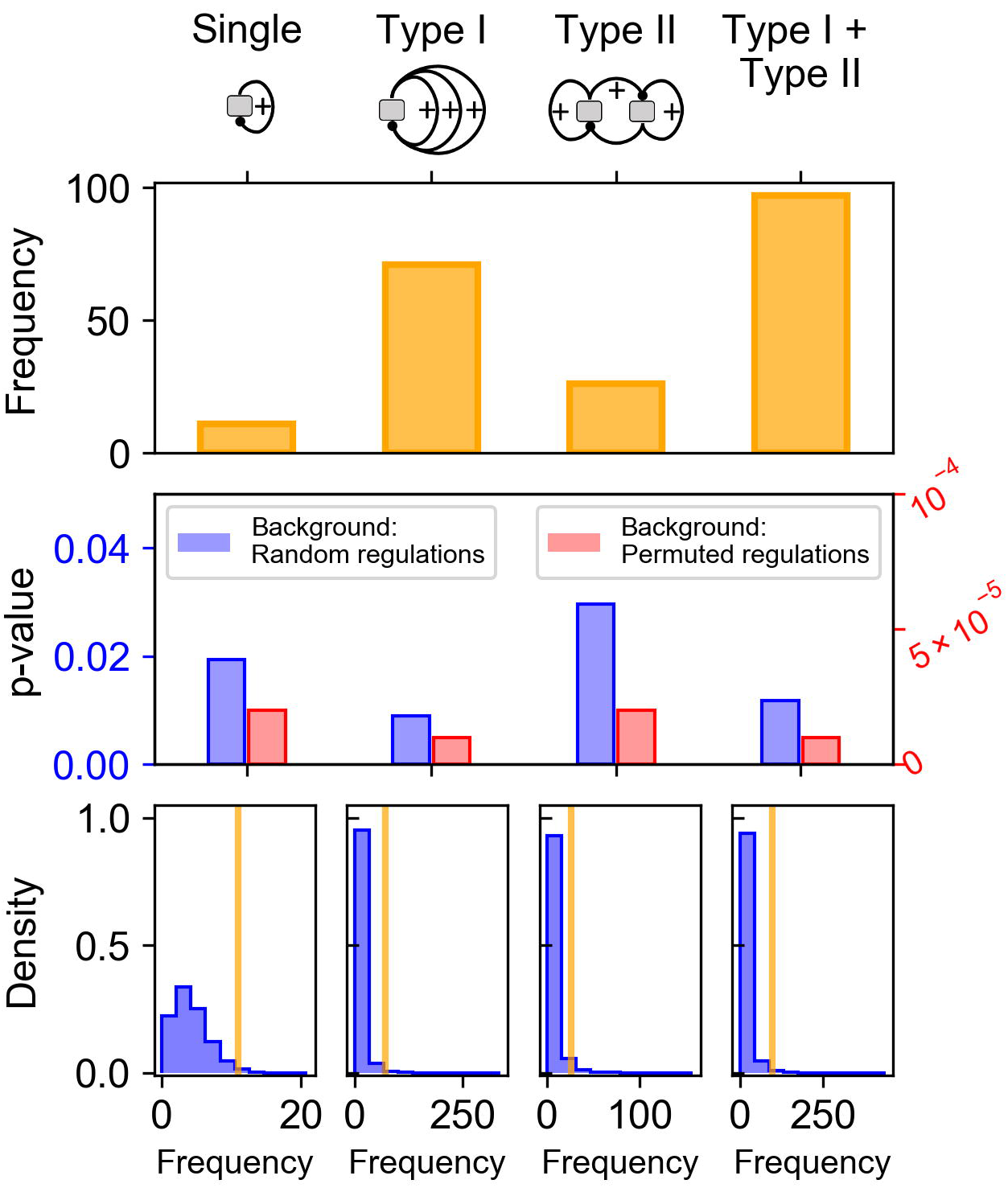
Enrichment of Type I and Type II motifs in the T cell model. Top panel: total occurrences of various types of motifs in the T cell network. Middle panel: empirical p-values of the single positive feedback loops and the sum of the two types of motifs. Bottom panel: an illustration of the p-values with the distributions of background population. Random networks were obtained by 1) permuting the regulations in the existing network by randomly assigning their sources and targets (red) and 2) assigning random regulations (positive, negative or none) between each pair of TFs (blue). 10^5^ random networks were generated with each method. Empirical p-values were obtained by counting the number of the random networks with the number of motifs not less than those in the T cell network. See Methods for details of the p-value definition. Distributions of motif frequencies obtained from the random networks using the second method are shown in the bottom panel. The yellow vertical bars represent the number of occurrences in the T cell network. The right-tail areas defined by the vertical bars correspond to the p-values shown in the middle panel (blue bars).

Since the minimum motifs alone can generate the four-attractor system, we asked whether the combination of these motifs enhances the ability of the network to produce the system. We therefore compared a subnetwork containing only one minimum Type I motif with another one containing multiple such motifs in terms of the performance to generate a particular four-attractor system (Figure 5A. See Methods and Text S1 for details). We found that the subnetwork with multiple Type I motifs outperforms the one with only one motif (Figure 5B and C, the purple curve for production rate has a more robust pattern showing 7 intersections with the degradation curve than the red production curve does), suggesting the advantage of combining multiple motifs with similar functions to enhance its overall performance. We next asked whether the topologies that contain both Type I and Type II motifs have greater probabilities to generate the four-attractor system than the topologies with one type of motifs do. When we explored the parameter space randomly for each topology with a fixed number of samples, a larger number of parameter sets that can generate the four-attractor system were found with the topologies containing both motifs than with those containing either Type I or Type II motifs only (Figure S5 and Figure 5D). This suggests that the combination of both motifs might be a robust strategy to generate the four-attractor system. This pattern was observed for all the topologies in the complexity atlas (Figure S6) as well as those with the same degree of complexity (Figure 5D, networks with 7 regulations were chosen because they have comparable fractions of the three types of motifs).

**Figure 5.**
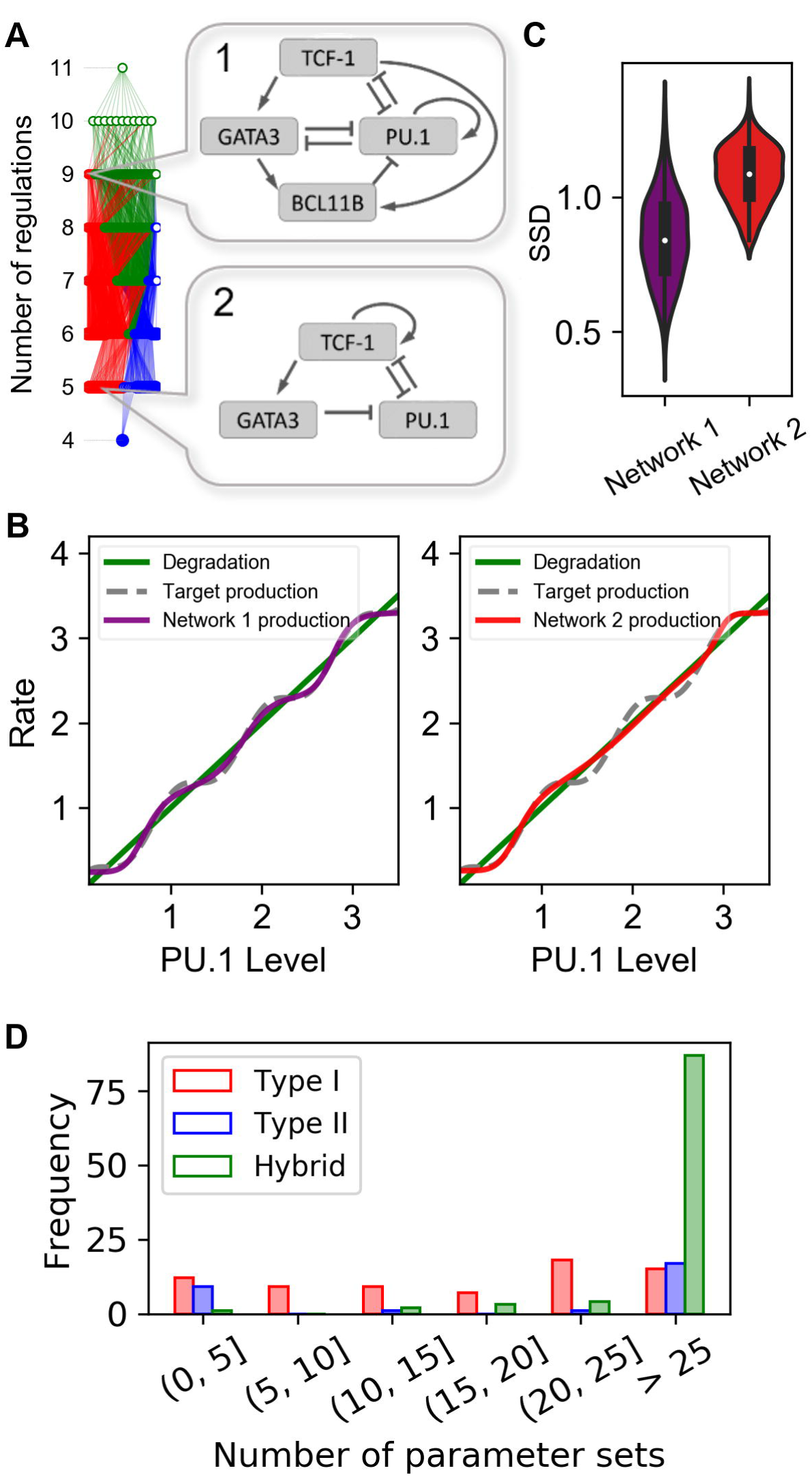
Comparisons of motifs with different complexity and types. **A.** Two specific network topologies were selected for comparing models with different complexity. Network 1 contains multiple Type I motifs, whereas Network 2 is a single Type I motif. The color code of the complexity atlas is the same as that in Figure 2 and Figure 3. Red: Type I motif. Blue: Type II motif. Green: Hybrid motif. **B.** Performances of the two subnetworks are compared. Performance was quantified with the sum of squared distance (SSD) from a predefined continuous production function (gray curve) of PU.1 level that generate four attractors (see details in supplementary text). Purple and red curves represent the optimized functions fitted to the gray curve. **C.** SSD values obtained from 500 optimization runs. Each value was calculated using the procedure shown in B. **D.** Histogram for the numbers of topologies with 7 regulations with respective to the number of parameter sets that generate the four-attractor systems per 10^6^ random parameter sets.

In summary, we found that the core transcriptional network controlling early T cell differentiation are enriched with Type I and Type II network motifs. The network composed of these two types of motifs governs a dynamical system containing four attractors, corresponding to four known stages in the early T cell development. The networks with both types of motifs and greater number of such motifs have more robust capability of generating the four-attractor systems than those networks with fewer types of numbers of motifs do.

### Stepwise transitions with restricted reversibility provide robustness to fluctuating differentiation signal to multiple intermediate states

We next characterized the dynamical features of the four-attractor system of the T cell development model in response to differentiation signals. For this and subsequent analysis, we focused on a model describing the network topology shown in Figure 3A (the full model). We first performed bifurcation analysis of the system to the changes of Notch signaling (Figure 6A). With the increasing Notch signal, the system undergoes three saddle-node bifurcations, at which the stability of the proceeding cellular states is lost (Figure 6A, black arrows). These bifurcation points therefore represent the cell state transitions from one stage to the next. The structure of the bifurcation diagram shows a remarkable robust multistep commitment program governed by the T cell transcription network: the commitment to each stage of the program has restricted reversibility in that the attenuation or withdrawal of the Notch signaling does not result in de-differentiation of the developing T cells (i.e. the return of the transcription profile to earlier stages that may have greater multipotency). It was previously shown that the commitment from DN2a to DN2b is an irreversible process with respect to Notch signaling, and this transition eliminates developing T cells’ potential to be diverted to any other lineages when Notch signaling is abolished [20,41]. However, simple toggle-switch models do not explain the observation that the multipotency of the early T cells is lost in a stepwise manner. For example, cells at ETP can be differentiated into B cells, macrophages, dendritic cells (DCs), granulocytes, natural killer (NK) cells and Innate lymphoid cellsubset2 (ILC2), whereas the potentials to commit to many of the lineages are blocked even in the absence of Notch signaling at the DN2a stage, at which the cells can only be differentiated into NK cells and ILC2 [20]. Therefore, the stepwise, irreversible transcriptional transitions revealed by our model is consistent with the experimental observations with respect to the loss of multipotency in the stepwise manner.

**Figure 6.**
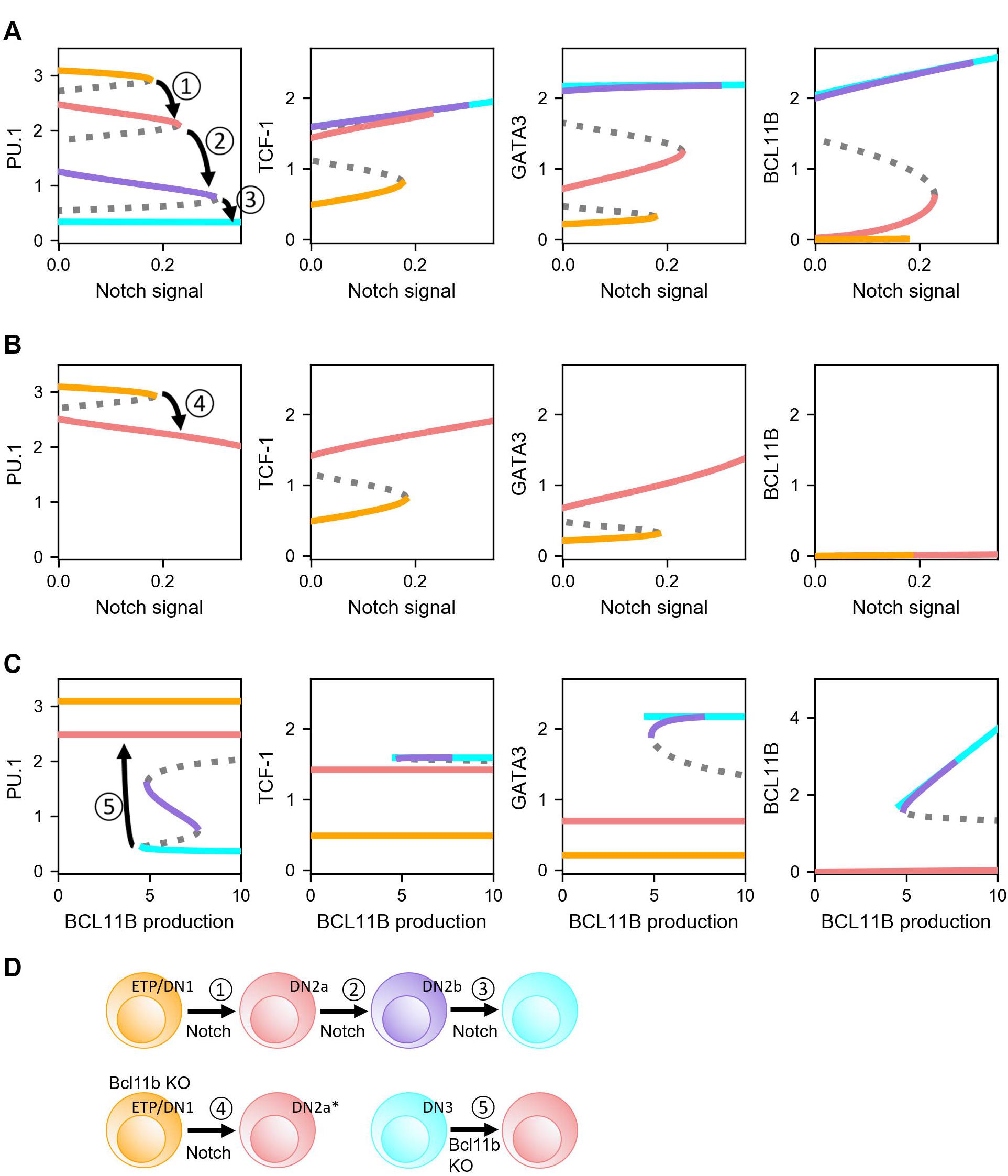
Stability analysis of the T cell model. The full model shown in Figure 3A is used for all the analysis.**A.** Bifurcation diagrams for the steady states of the four core factors with respect to the Notch signal. Solid curve: stable steady state. Dashed curve: unstable steady state. **B.** Bifurcation diagram under Bcl11b knockout condition with respect to Notch signal. Solid curve: stable steady state. Dashed curve: unstable steady state. **C.** Bifurcation diagram with respect to BCL11B production rate parameter. Solid curve: stable steady state. Dashed curve: unstable steady state. **D.** Illustration of the observed transitions among the four states. Colors of the stable branches of the bifurcation diagrams and the cell icons are matched to the cellular states shown in Figure 1.

Although the absence of Notch signal does not allow the reversal of lineage progression, it was previously shown that the absence of BLC11B in lymphoid progenitor cells blocks its ability to progress to DN2b stage, whereas the Cre-controlled knockout of BCL11B in committed T cells (e.g. DN3 cells) reverts its transcriptional profile to DN2a-like cells [28]. Upon blocking the production of BCL11B in our model, we observed the loss of attractors of DN2b and DN3, and the DN2a state is the only stable stage even in the presence of the strong Notch signaling (Figure 6B). As a result, increasing Notch signaling only triggers one saddle-node bifurcation, representing the transition from ETP to DN2a cell (Figure 6B, top panel and black arrow), whereas the decrease of the BCL11B production triggers the transition back to DN2a instead of ETP (Figure 6C). These results are in agreement with the previous experimental findings [28], and they further support the importance of the multistep differentiation system revealed by our model.

The bifurcation analysis shows how the lineage progression is influenced by stably increasing or decreasing Notch signal strengths. We next asked how the duration of Notch signal may control the multistep lineage transition. By inducing the differentiation with varying durations of the Notch signaling, we found that cells experiencing transient Notch signals may only commit to intermediate stages of differentiation (Figure 7A). In addition, the system is able to integrate the information of the signal intensity and duration to make decision on the lineage progression. These results suggest that the multistep lineage transition can be triggered by the increasing strength of the signal, the increasing duration of the signal, or the combination of both types of signal dynamics. Earlier experimental studies have shown that transient Notch signaling can irreversibly drive the cells to an intermediate, but committed stage with a definitive T cell identity (DN2b) [28,41,43]. This is in agreement with our results, and our model further suggests that the commitment to other intermediate states is also irreversible with respect to the lineage progression (note that this irreversibility does not refer to the establishment of T cell identity).

**Figure 7.**
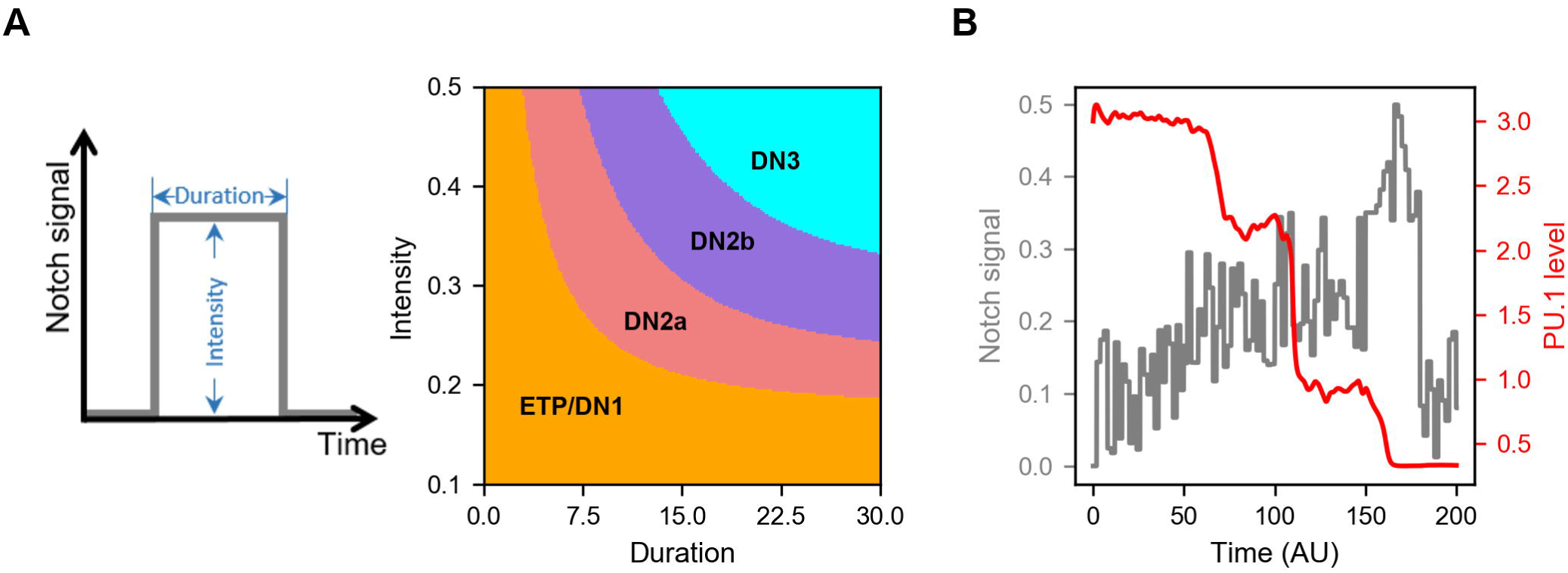
Multistep lineage transitions under the influence of varying dynamics Notch signals. **A.** Strength and duration of the Notch signal were varied in each simulation. 200X200 combinations of different signal strengths and durations were tested, and the final cellular phenotypes were determined using the levels of the four core factors. **B.** Dynamics of PU.1 in response to increasing Notch with significant fluctuations. The mean of the Notch signal increases linearly in the first phase, then it is attenuated in the second phase. Fluctuations were simulated with additive noise in small time intervals.

One possible advantage of the multi-stable system is its robustness of response in facing fluctuating signals. We therefore performed numerical simulations of the dynamical system under increasing Notch signaling with significant fluctuations. Under this condition, transient reduction of Notch signaling halted the progress of the lineage commitment but did not trigger the de-differentiation (Figure 7B). Our model suggests that the design of transcriptional network allows system to stop at intermediate stages before proceeding to the next ones. This strategy has several potential physiological benefits: 1) it protects the cell lineage progression against sporadic fluctuations of Notch signaling; 2) it facilitates the ‘checkpoints’ before lineage commitment in the middle of the entire developmental process and 3) it allows the stable storage of differentiation intermediates which can be differentiated into mature T cells rapidly when there is an urgent need of new T cells with a diverse T cell receptor repertoire.

### Quantitative analysis of the energy landscapes and minimum action paths delineates the patterns of the multiple-attractor system in T cell differentiation

With the deterministic modeling and bifurcation approaches, we described the local stability for multi-stable T cell model. However, the global stability is less clear from the bifurcation analysis alone. In addition, it is important to consider the stochastic dynamics for T cell development model, because the intracellular noise may play crucial roles in cellular behaviors [44,45]. The Waddington landscape has been proposed as a metaphor to explain the development and differentiation of cells [46]. Recently, the Waddington epigenetic landscape for the biological networks has been quantified and employed to investigate the stochastic dynamics of stem cell development and cancer [47-51].

Following a self-consistent approximation approach (see Methods), we calculated the steady state probability distribution and then obtained the energy landscape for the model of the early T cell development. For visualization, we selected two TFs (PU.1 and TCF-1) as the coordinates and projected the 4-dimensional landscape into a two-dimensional space, by integrating the other 2 TF variables. Here TCF-1 is a representative T cell lineage TF, and PU.1 is a TF for alternative cell fates. Note that our major conclusions do not depend on the specific choice of the coordinate (see Figures S7 and S8 for landscapes with PU.1/BCL11B and PU.1/GATA3 as the coordinates).

In the case without Notch signal (N = 0), four stable cell states emerge on the landscape for the T cell developmental system (Figure 8). On the landscape surface, the blue region represents lower potential or higher probability, and the yellow region represents higher potential or lower probability. The four basins of attraction on the landscape represent four different cell states characterized by different TF expression patterns in the 4-dimensional state space. These states separately correspond to ETP/DN1 (high PU.1/low TCF-1/low BCL11B/low GATA3 expression), DN3 state (low PU.1/high TCF-1/high BCL11B/high GATA3 expression), and two intermediate states (DN2a and DN2b, intermediate expression for the four TFs). The existence of four stable attractors is consistent with experiments [16-19]. As the Notch signal (N) increases, the landscape change from a quadristable (four stable states coexist), to a tristable (DN2a, DN2b and DN3), to a bistable (DN2b and DN3) and finally to a monostable DN3 state (Figure S9). These results provide a straightforward explanation for the irreversibility observed in experiments for the stepwise T cell lineage commitment.

**Figure 8.**
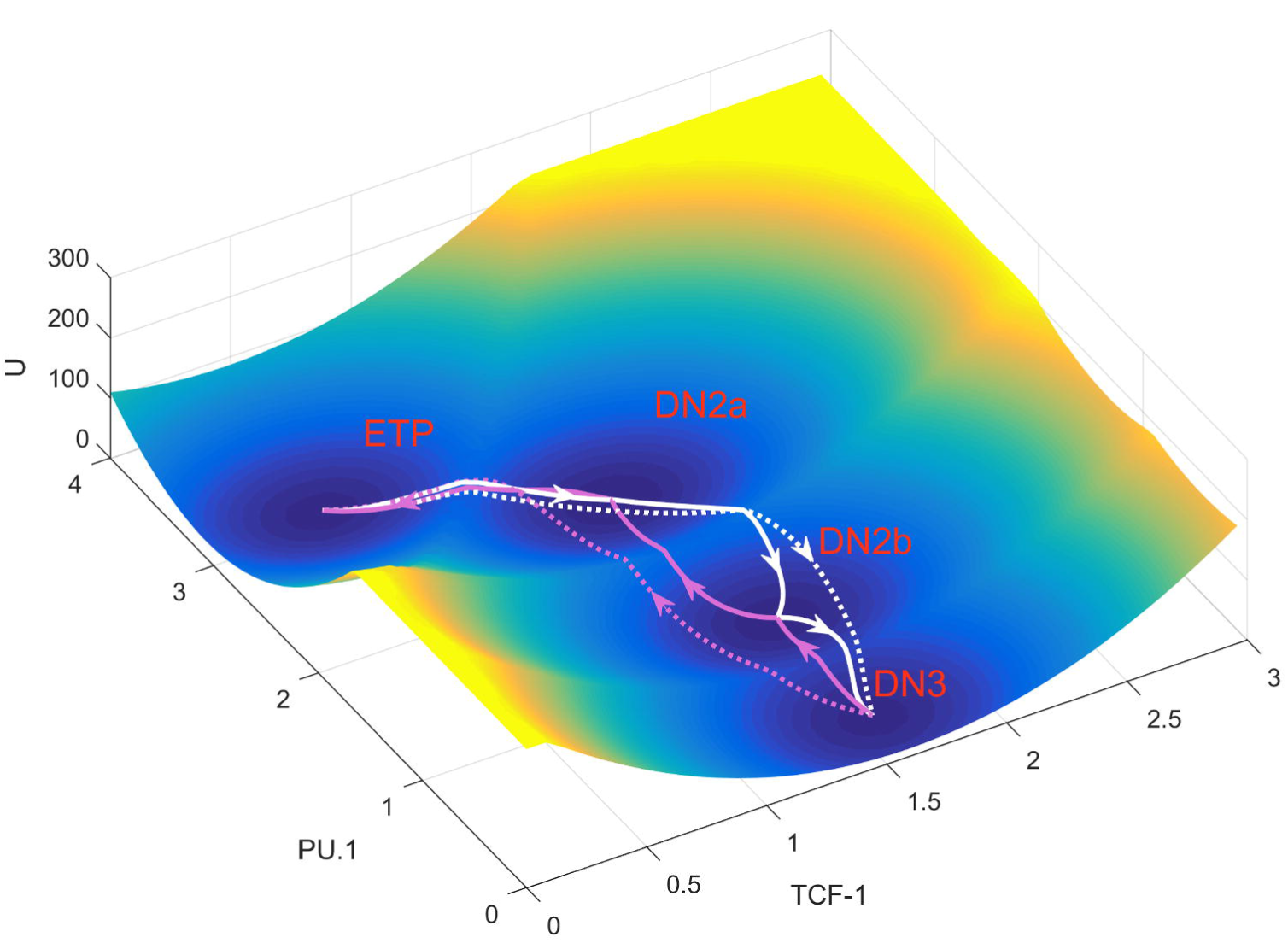
Energy landscape for T cell development. The landscape and corresponding minimum action paths (MAPs) for the T cell developmental network are shown in 3-dimensional figure. White solid lines represent the MAP from ETP state to DN2a, DN2b, and DN3 states. Magenta solid lines represent the MAP from DN3 to DN2b, DN2a, and to ETP state. Dashed lines represent the direct MAP from ETP to DN3 and from DN3 to ETP states, respectively. Here, TCF-1 and PU1 are selected as the two coordinates for landscape visualization. See Supporting information for the landscapes using other pairs of TFs.

To examine the transitions among individual cell types, we calculated kinetic transition paths by minimizing the transition actions between attractors [52,53], obtaining minimum action paths (MAPs). The MAPs for different transitions are indicated on the landscape (Figure 8). The white MAPs from the ETP state to the DN3 state, correspond to the T cell developmental process while the magenta MAPs from the DN3 state to the ETP state, correspond to reprogramming process. The lines represent the MAPs, and the arrows denote the directions of the transitions. The MAP for T cell developmental process and the MAP for the backward process are irreversible, since the forward and reverse kinetic paths are not identical. This irreversibility of kinetic transition paths is caused by the non-gradient force, i.e. the curl flux [54,55]. Here, the solid white lines represent three stepwise transitions from ETP to DN2a, DNa2 to DN2b, and DN2b to DN3, whereas the dashed white line represents the direct transition paths from ETP to DN3. From the MAPs for T cell development, we found that the direct transition path is very similar to the stepwise transition path (the white solid line is similar to the white dashed line, Figure7, Figures S7 and S8), which indicates that the T cell developmental process needs to go through the two intermediate states (DN2a and DN2b). This confirms the critical roles of the intermediate states for the T cell differentiation. It is worth noting that the MAPs here quantify the most probable transition paths, which suggest the optimal path (with least transition actions) for cells to switch from one state to another. However, in a realistic gene regulatory system, usually a signal is needed to induce cell state transitions (e.g. the Notch signaling is used here to induce T cell development).

To investigate the dynamical developmental process of T cell for multiple TFs, we visualized the 4-dimensional MAP from the ETP to the DN3 state by discretizing the levels of the four TFs. We found that for T cell development, TCF-1 is upregulated first, followed by the activation of GATA3. This leads to the complete inactivation of the alternative fate TF PU.1 and the activation of BCL11B (Figure 9). Interestingly, this temporal order is in good agreement with experimental observations [56]. These results suggest that the sequence of switching on or off for different TFs can be critical for the lineage commitment of T cell development. Moreover, under the Bcl11b knockout condition (kB=0), the landscape changes from a quadristable (four stable states coexist), to a bistable (ETP and DN2a) state (Figure S10), which is consistent with the bifurcation analysis (Figure 6) and experimental observations [28].

**Figure 9.**
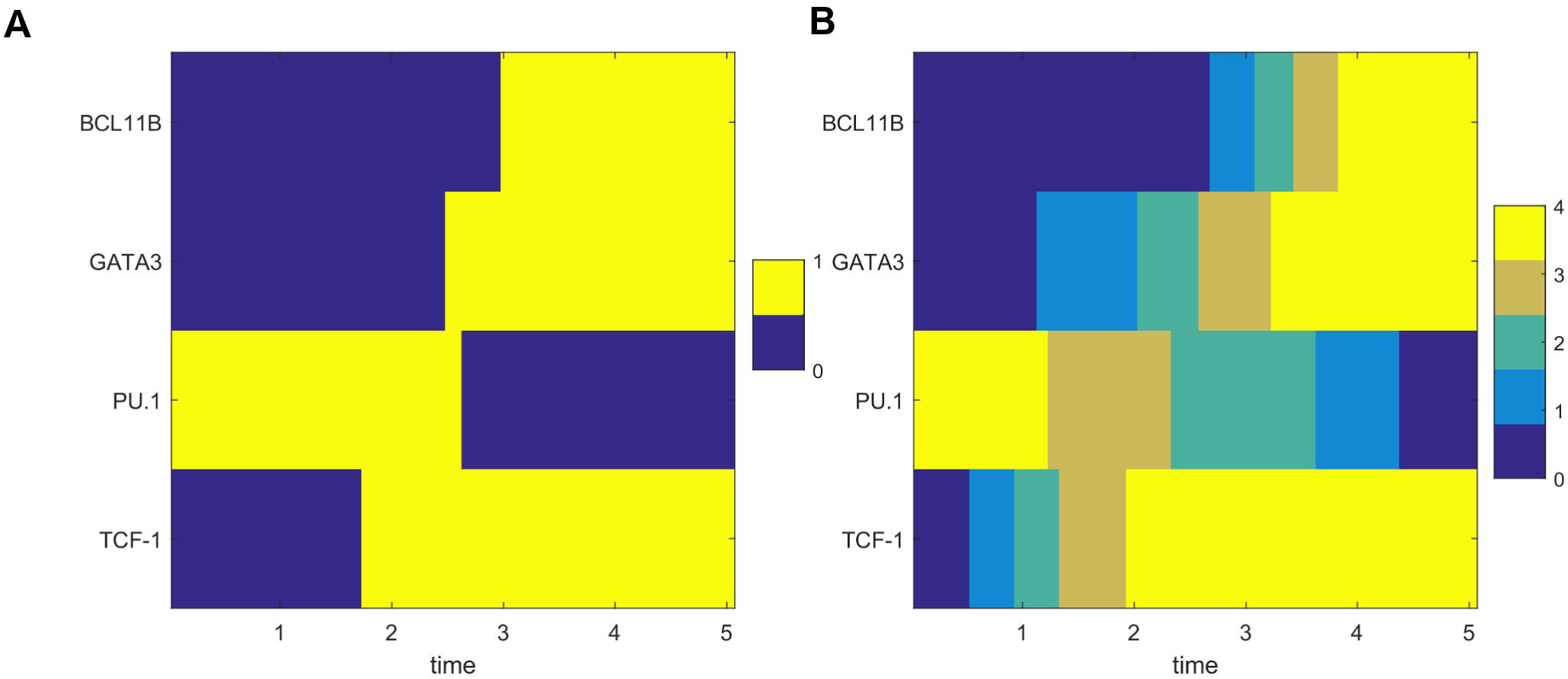
Discrete kinetic transition paths for T cell model. Transition paths from ETP state to DN3 state in terms of levels of 4 different TFs. **A.** The relative TF levels are discretized to 0 or 1. 1 represents that the corresponding TFs are in the on (activated) state and 0 represents that the corresponding TFs are in the off (repressed) state. **B.** The relative TF levels are discretized to five values from low to high. X axis shows the time along the transition path.

### Global sensitivity analysis based on landscape topography reveals the critical factors for T cell development

To identify the critical factors (regulations and TFs) which determine T cell development, we performed a global sensitivity analysis based on the landscape topography. Specifically, we use the transition action between attractors as a measure to quantify the feasibility of a transition between different attractors. A smaller transition action, corresponding to a larger energy barrier, means a more feasible transition from one attractor to another. In this way, by changing the parameters each at a time we can identify the critical parameters for T cell development (we use the transition from ETP to DN3 as an example). To do this, we constrict the models within the parameter region corresponding to the four-attractor system, so that we can make comparisons for the changes of transition actions as parameters are varied.

We identified some critical parameters of which the variations caused significant changes of transition actions between ETP and DN3 attractor. These parameters include the effective degradation rate of PU.1, (rdP), the regulated production rate of PU.1 (kP), the basal production rate of PU.1 (kP0), the threshold of the self-activation of PU.1 (KPP), and the threshold for the repression of PU.1 on GATA3 (KGP) (Figure 10). In particular, the increase of the self-activation strength of PU.1 (i.e. decreased KPP) reduces the transition action from DN3 to DN2b (Figure 10B), indicating a less stable DN3 state and a more stable ETP state. This is reasonable because the PU.1 is a major TF for alternative cell fates (B-cell, dendritic-cell, and myeloid cell), and silencing of PU.1 is operationally important for T cell commitment [28]. Additionally, the increase of the repression strength of PU.1 on GATA3 (decreased KGP) raises the transition action from ETP to DN2a (Figure 10B), indicating a more stable ETP state and a less stable DN3 state, which is consistent with the observation that GATA3 is a critical TF promoting T cell development. Overall, these results from sensitivity analysis indicate that the PU. 1 synthesis/degradation related parameters, the GATA3 synthesis related parameters, and the regulations between PU.1 and GATA3 are critical to the dynamics and the cell fate decisions of T cell development. This indicates that the regulatory circuit between PU.1 and GATA3 plays critical roles for the cell fate determinations during T cell development.

**Figure 10.**
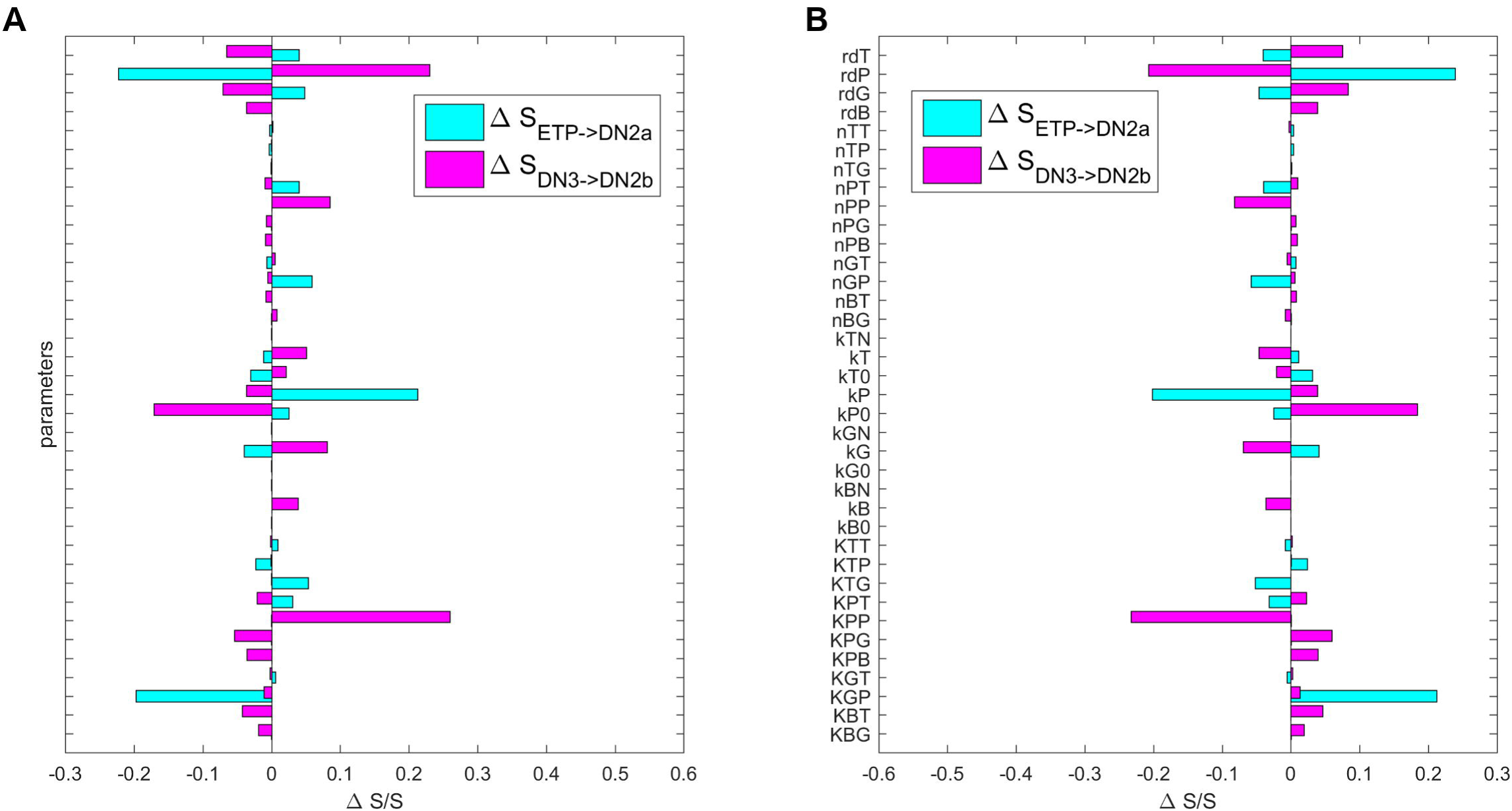
Global sensitivity analysis for T cell developmental model. Sensitivity analysis was performed for the 39 parameters in the T cell model. The transition actions between different states (S_ETP→DN2a_ and S_DN3→DN2b_) were calculated to quantify the sensitivity of parameters on the landscape. The Y-Axis represents the 39 parameters. The X-Axis represents the percentage of the transition action (S) changed relative to S without parameter changes. Here, S_ETP→DN2a_ represents the transition action from attractor ETP to attractor DN2a (cyan bars), and S_DN3→DN2b_ represents the transition action from attractor DN3 to attractor DN2b (magenta bars). A. Each parameter is increased by 1%, individually. B. Each parameter is decreased by 1%, individually.

## Discussion

In this study, we identified two types of network motif families that are responsible for generating a four-attractor dynamical system commonly observed in stepwise cell differentiation. Some instances of these motifs were previously described and analyzed in the context of binary or ternary switches during lineage transitions [57-61], but the systematic analysis for these motifs was not performed to our knowledge. In addition, the design principle for multiple intermediate states was not clear. Our approach provides a comprehensive framework for analyzing systems with a complex dynamical property, a four-attractor system with stepwise transcriptional modulation, and we illustrate the intricate relationships among these motifs with an intuitive visualization method.

Previous studies on biological circuits governing irreversible transitions focused on the analysis of toggle switches which generate none-or-all type of responses [62,63]. Our work suggests that multistep or graded responses can be associated with irreversible transitions as well. Given the importance of graded response in various biological scenarios [64-66], we expect the design strategy that we found can be useful for discovery of natural-occurring irreversible graded responses or construction of synthetic biological circuits producing these responses. Our work also suggests that the response to signals, or the progression of lineage transition, may be proportional to the intensity and/or the duration of the signal. This is consistent with the previous observations that the duration of the morphogen signal can be critical for cell lineage choice [67,68]. Of note, when signal strength is converted to digital (none-or-all) response in early phases of signal transduction, its duration can play an essential role in determining the graded response [69].

In our systematic exploration in the network topology space, we took the assumption that network structure is correlated with its function, i.e. assuming the existence of functional motif structure in transcription regulatory networks. The notion of network motifs is very helpful for understanding many complex biological systems [70,71], but the richness of dynamic behaviors of these motifs is beyond their structures – distinct kinetic rates in the same motif can produce diverse responses [72]. Therefore, it is expected that the motifs that we discovered may be able to generate dynamical behaviors different from the four-attractor system (we will discuss some of them in the following paragraphs). We also expect that some of network motifs can be responsible for multiple functions by themselves, and this multifunctionality may explain the diverse motifs that we found for the four-attractor systems in the biological examples. Future work is warranted to examine the distributions of the diverse functions in the parameter space of the motifs that we found. Nonetheless, it is important to understand the capacity of the network motifs in terms of their functional outputs. Our work provides a holistic view of the potential network motif structures governing multistep cell lineage transitions.

Although network motifs with three positive feedback loops closing at a single factor (Type I motifs discussed in this study) were not systematically analyzed in previously studies to our knowledge, some simpler versions of Type I motif, e.g. a pair of interconnected positive feedback loops, have been described in various systems such as the epithelial-mesenchymal transition and the cancer progression [59,73]. These systems typically govern ternary switches with a single intermediate state. These studies and ours suggest a correlation between the number of positive feedback loops and the number of the intermediate states the system may be able to generate. In fact, early studies on multistability systems have shown the requirement of positive feedback loops for generating multiple steady states [74], which was later proved mathematically [75]. Intriguingly, an ultrahigh feedback system similar to the Type I motifs was shown to govern irreversible transitions with low differentiation rates for adipocytes [76]. It would be interesting to examine whether controlling the low differentiation rate through cell-to-cell variability and controlling the number of intermediate states suggested by our model can be achieved in the same system. Our findings are consistent with the earlier work in that they highlight the importance of this type of signaling motifs in controlling cell differentiation by preventing the direct and homogeneous transition from the initial state to the final one.

Near symmetrical parameters in models based on a particular instance of the Type II motif class (the one with mutually inhibiting TFs) have been widely used to explain stochastic lineage choice observed in embryonic stem cells, developing hematopoietic cells and CD4^+^ T cells [77,78]. Our findings with Type II motifs complement these studies with newly identified functions of these motifs for cell differentiation. Instead of the stochasticity that breaks the symmetry of this motif, the Notch signal may be responsible for switching the system from one side (PU.1 high) to another (PU.1 low) in a stepwise fashion, and the intermediate states mark the stable stages where the system is relatively balanced in terms of two groups of competing TFs.

It was previously suggested that the network consisting of four core transcription factors governs a bistable switch with irreversible transition [41]. Our models based on this network provide explanations for additional experimental observations with respect to the multistep feature of the early T cell development. Although it is possible that interconnection of multiple positive feedback loops simply enhances the robustness of the bistable switches, the observation that several important irreversible transitions in cell cycle progression are primarily controlled by two positive feedback loops implies that the enrichment of the positive feedback loops in the T cell transcriptional network is unlikely due to the intrinsic biophysical limits of positive feedback loops in generating bistable switches [63,79]. Instead, other cellular functions, such as generating the multiple intermediate states, might be the performance objectives for the design of this network.

Our model of early T cell development suggests that the differentiation program may be stopped at multiple locations in the state space of transcription levels of key factors. These multiple attractors may correspond to the lineage branching points at which the progenitor cells are given opportunities to be converted to T cell as well as other types of lymphocytes. As such, it is possible that this dynamical property is exploited to achieve a better control for the fate determination of the lymphoid progenitor cells at systems level. Given that subpopulations of NK cells and DCs are generated by the thymus [80-82], the multistep lineage transition provides a basis for channeling the lymphoid progenitor to multiple lineages in a precise manner.

Based on the recent landscape-path theory and the T cell gene regulatory network model, we investigated the stochastic dynamics of T cell development. We identified four stable cell states characterized by attractors on the landscape including ETP/DN1, DN3, and two intermediate states (DN2a and DN2b). We also calculated the kinetic transition paths between different cell states from minimum action path approaches. Importantly, from the MAPs of T cell development, we found that different TFs are switched on or off in different orders. For example, TCF-1 needs to be first activated, and then GATA3 is activated, leading to the inactivation of PU.1 and activation of BCL11B. These predictions agree well with experiments [28,56], which provides further validations for our mathematical model.

In our models, we only considered four core factors based on previous published T cell gene regulatory network for simplicity [41]. In the realistic biological system, there are more factors critical to T cell development [28]. It would be interesting to incorporate other important factors into the network and construct a more realistic model for T cell development. By studying the landscape of more comprehensive T cell development network, we will better understand the underlying regulatory machinery and obtain more insights into the intricate mechanisms for T cell development.

In summary, we identified a large family of network motifs that can generate four attractors that are observed in various biological systems involving cell lineage transition. We built a mathematical model for transcriptional network controlling early T cell development, and we found that the network underlying this developmental process is enriched with the motifs that we identified. The system with the four attractors has a remarkable irreversibility for transitions to multiple intermediate states when the differentiation signal is varied. We suggest that this multistep process may be useful for precise control of the differentiation of lymphoid progenitor cells towards T cell and other cell types. Our T cell model provides new insights into the complex developmental or regeneration processes, and our combined approaches of comprehensive analysis of network motifs for generating multistable systems and landscape-path framework provide a powerful tool for studying a wide range of networks controlling cell lineage transitions.

## Methods

### Framework of mathematical modeling

We used ordinary differential equations (ODEs) to describe the dynamics of the concentrations of transcription factors (TFs). We used Hill function to describe the transcriptional regulation by TFs. Each ODE has the following form:

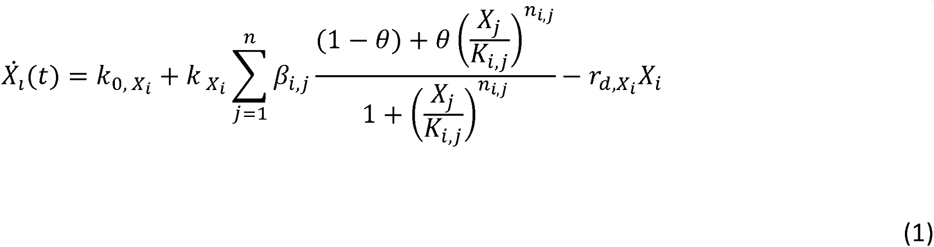

Here,*X*_*i*_ represents the concentration of a transcription factor (TF). *k*_0,*x*_*i*__ is the basal production rate of the TF in the absence of any regulator. *k*_*x*_*i*__. is the maximum production rate under the control of the transcriptional activators and inhibitors of this TF. *β*_*tj*_ denotes the weight of the influence of the TF *j*on *i*.The sum of the Hill functions determines the regulation of the production of this TF by other TFs. In each term of the summation, *θ* = 1 when the regulating TF (*X*_*j*_) is an activator. *θ* = 0 when the regulating TF is an inhibitor. *K*_*i,j*_ is the apparent dissociation constant of the regulating TF binding to its regulatory element of the promoter, and it describes the effectiveness of the regulation in terms of the concentration of the TF. *n* is the total number of regulating TFs. *r*_*d,x*_. is the effective degradation rate constant. The production rate of the proteins is assumed to be linearly correlated with mRNA production rate. Similar generalized forms of Hill function were previously used for analysis of a variety of gene regulatory networks [48,83]. One time unit of our model corresponds to 20 minutes, and all the parameters are dimensionless.

To exclude the possibility that our conclusions are sensitive to the choice of the form of equations, we used an alternative form of ODE to describe the regulatory networks:

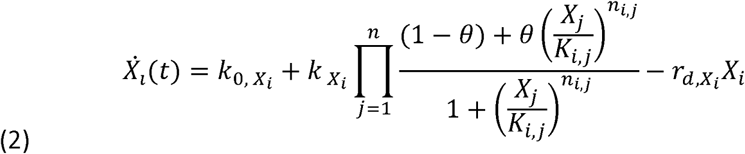

In these ODEs, multiplication of Hill functions was used instead of addition. Similar forms of Hill function were also previously used for modeling a variety of gene regulatory networks [60,84]. With this form, the two types of network motifs that generated the four-attractor behavior are the same as those discovered with the additive form of Hill functions (Figure S3). In fact, using both forms of equations gave rise to the same number of network topologies (216 topologies with the steady states shown in both Figure S3 and Figure S4). Therefore, our conclusions are robust in terms of the choice of equation form.

During topology searching, random parameters values were chosen from defined ranges (Table S1, see below).

### Topology searching for four-attractor systems

Network topology searching was first performed for all possible topologies involving up to 3 nodes (TFs) and 6 regulations that are able to generate four-attractor systems with stepwise changes of TF levels.

Three-node networks were previously used to explore several types of functional dynamics of network motifs [38,85]. Isometric topologies were removed in the search. For each topology, we performed random sampling with 10^6^ parameter sets. For each parameter set, we selected 125 initial conditions in the three-dimensional state space ((0, 3.3) for each variable) using Latin Hypercube sampling, and then solved the ODEs numerically. We stopped the simulations at time point 500 and checked if the 125 ODE systems are stabilized at four or more distinct steady states. We next checked if the changes of the TFs are monotonically coupled. We first ordered the steady states by the levels of one TF, and then we looked for scenarios in which all other TFs monotonically increase or decrease with the ordered TF (i.e. the attractors with stepwise changes of the TFs). We excluded the scenarios in which one TF is not monotonically correlated with others in terms of their levels at the four attractors. Models that generated oscillations at the final time point were also excluded. The parameter sets which produced the stepwise changes of steady state were accepted and their associated network topologies were analyzed. Parameter values for the minimum topologies are listed in Table S2.

Complexity atlas was plotted for the obtained network topologies as described previously by Jiménez et al [40] (Figure 2B and Figure 3C).

### Transcriptional network model for early T cell development

We built a model for early T cell development based on the regulations that were previously shown experimentally [86-99]. Information about experimental evidence is described in Table S3. The form of equations is similar to Equation (2). We chose this multiplicative form of Hill functions because earlier experimental study suggested that regulations of Bcl11b gene are combined via an ‘and’ logic gate [100], which favors the use of multiplication. Although similar detailed information is not available for other TFs, we have shown that our main conclusions with respect to the multistep transitions controlled by a network motif family do not depend on the choice of the form of equations (Figure 2 and Figure S4). Full list of equations is included in Text S1. The parameter values were obtained by random searching described above followed by minor manual adjustment. The parameter values are listed in Table S4. To explore the subnetworks of the T cell development model that are essential for the four-stage transition, we performed similar exhaustive search in a set of 1553 non-redundant topologies (2047 subnetworks) to find functional circuit in the model. We obtained 568 topologies (701 topologies) from the search, and we analyzed them with complexity atlas. Isometric topologies were removed in the simulations, but they are included in the complexity atlas so that we do not mix isometric topologies with possibly differential biological meanings specific to certain genes.

During bifurcation analysis, the value of the parameter *N* (Notch signal strength) or *k*_BCL11B_ (maximum production rate of BCL11B) is varied and the changes of the steady states of the system were analyzed. We let *k*_BCL11B_ = 0 to simulate the *Bcl11b* knockout condition.

To simulate the system under various scenarios of Notch signaling, we first varied the strength and/or duration of the Notch signal and checked the steady state distribution of the system under the varying strengths and durations. We tested 200X200 combinations of strengths and durations of Notch signals and obtained the phenotypes of the cells at the steady state. To simulate the fluctuating Notch signals, we divided the time window of the simulation into small intervals (0.1 unites of time). For each interval, we used a random number with a specified mean and an additive noise. The mean of the Notch signal first increased overtime and then became attenuated.

### Enrichment analysis of the four-attractor motifs in the T cell model

To quantify the enrichment of various types of motifs, we used the generic definition of p-value: the p-value for a particular motif is the probability of obtaining at least n number of motifs from a random network population, where *n* is the observed number of such motif in the T cell network. To compute the p-values, we first counted the frequencies of the positive feedback loop, Type I motif and Type II motif in the T cell model (i.e. *n*_l_, *n*_2_,*n*_3_, *n*_4_ representing the numbers of positive feedback loops, Type I motifs, Type II motifs, and the sum of the Type I and Type II motifs respectively). Random networks were generated using two methods: 1) for each regulation in the existing T cell model, we randomly reassign its source and target TFs (referred to as ‘permuted regulations’), and 2) for each pair of TFs from the network, we randomly assign a regulation (positive, negative or none) (referred to as ‘permuted regulations’). For each of the two methods, we generated 10^5^ networks, and we calculated the empirical p-values by counting the number of the random networks with the numbers of motifs not less than those of respective motifs in the T cell network. The method with permuted regulations is more biologically relevant because the number of the positive and negative regulations are retained in the random networks. We used the second approach as alternative to exclude the possibility that the conclusion of the trend of the p-values is due to the low number of networks containing the extreme amount of the motifs.

### Optimization for performance comparison of two subnetworks

Due to the difficulty to compare the performances of regulatory circuits with different complexities in general, we selected two specific instances of Type I network motif for comparison. One of them contains only one Type I motif, whereas the other one contains multiple motifs. For each topology, we reduced the system to one ODE with quasi-steady state assumption and defined a continuous production rate function that can produce four attractors as a surrogate function (see Text S1). Multiple runs of optimization using differential evolution algorithm was used, and 500 converged parameter sets for each circuit were used for comparison. This optimization method was previously used for finding optimum parameter sets and for comparing the performances of regulatory circuits [58,101,102].

### Self-consistent mean field approximation for the quantification of energy landscape

The temporal evolution a dynamical system was determined by a probabilistic diffusion equation (Fokker-Planck equation). Given the system state P{*X*_1_,*X*_2_,…,*X*_*N*_t), where*X*_1_,*X*_2_, …, *X*_*N*_ represent the concentrations of molecules or gene expression levels, we have N-dimensional partial differential equation, which are difficult to solve because the system has a very large state space. Following a self-consistent mean field approach [48,54,103,104], we split the probability into the products of the individual probabilities: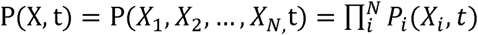 and solve the probability self-consistently. In this way, we effectively reduced the dimensionality of the system from MN to MN (M is the number of possible states that each gene could have), and thus made the computation of the high-dimensional probability distribution tractable.

Based on the diffusion equations, when the diffusion coefficient D is small, the moment equations can be approximated to [105,106]:

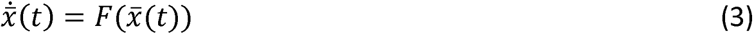

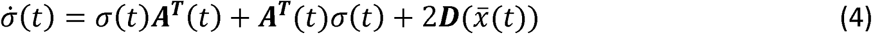

Here, 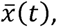 *σ*(*t*) and.***A***(*t*) are vectors and tensors. ***A***(*t*) denotes the covariance matrix and ***A***(*t*) is the jacobian matrix 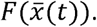 ***A***^*T*^(*t*) is the transpose of.*A*(*t*). The elements of matrix A are specified as: 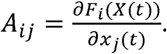 By solving these equations, we can acquire 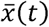 and *σ*(*t*). Here, we consider only the diagonal elements of u(t) from the mean field approximation. Then, the evolution of the probability distribution for each variable can be acquired from the Gaussian approximation:

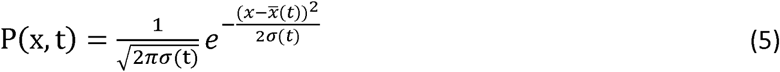

The probability distribution acquired above corresponds to one stable steady state or the basin of attraction. If the system has multiple stable steady states, there should be several probability distributions localized at each basin with different variances. Thus, the total probability is the sum of all these probability distributions with different weights. From the self-consistent approximation, we can extend this formulation to the multi-dimensional case by assuming that the total probability is the product of each individual probability for each variable. Finally, with the total probability, we can construct the potential landscape by: U(x) =—ln*P*_*ss*_(*x*). In this work, we define two quantities based on the landscape theory. One is the energy barrier height, which is defined as the energy difference between the local minimum and the corresponding saddle point. Another quantity is the transition action, which is defined as the minimum action from one attractor to the other. These two quantities both measure the difficulty of the transitions. However, the transition actions are suggested to provide a more accurate description for the barrier crossing between attractors or the transition rate {Feng, 2014 #113}. Therefore, we used the transition actions to quantify the difficulty of the transitions between attractors in this work (see the following section for minimum action paths).

### Minimum action paths from optimization

Following the approaches based on the Freidlin-Wentzell theory [52,107,108], for a dynamical system with multistability the most probable transition path from one attractor *i* at time 0 to attractor; at time T, 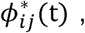 t ∈[0, T], can be acquired by minimizing the action functional over all possible paths:

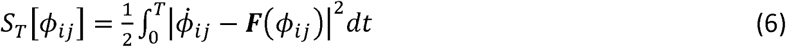

Here ***F***(*ϕ*_*jy*_) is the driving force. This optimal path is called minimized action path (MAP). We calculated MAPs numerically by applying minimum action methods used in [52,107].

## Supporting information

**Text S1. Model equations, model reductions and evaluation procedure through optimization.**

**Table S1. Ranges of parameter values for sampling 3-node networks.**

**Table S2. Parameter values for models of 29 minimum motifs for 3-node networks.**

**Table S3. Experimental evidence supporting the regulations in the early T cell development model.**

**Table S4. Parameter values for early T cell development model.**

**Figure S1. Phase planes for Type I minimum network topologies.** Nullclines for TF A (the node on the left of the network diagram) and TF B (the node on the right of the network diagram) are shown. Stable steady states are shown as black dots. The inset network diagram shows the corresponding network. Random parameter sampling was used to obtain the parameter sets that allows the 4-attractor systems.

**Figure S2. Phase planes for Type II minimum network topologies.** Nullclines for TF A (the node on the left of the network diagram) and TF B (the node on the right of the network diagram) are shown. Stable steady states are shown as black dots. Random parameter sampling was used to obtain the parameter sets that allows the 4-attractor systems.

**Figure S3. Overlaid four attractors for each of the 216 topologies from the 3-node network that produce 4-attractor systems.** Factor A denotes the TF on the left of the network diagram. Factor B denotes the TF on the right of the network diagram. In some topologies A and B and positively correlated (left panel), whereas they are negatively correlated in other topologies (right panel). Colored dots denote the stable steady states. Colored lines connect states of their corresponding topologies. The colors of the cell states match the illustration in Figure 1. The colors of the lines denote different representative models. z-score is calculated by shifting the mean of each four attractors to 0 and then normalizing the four data points to unit variance data. All models shown in this figure are built with additive form of Hill functions.

**Figure S4. Four-attractor systems generated with the alternative form of equations. A.** Overlaid four attractors for each of the 216 topologies from the 3-node network that produce 4-attractor systems. Factor A is the TF on the left of the network diagram. Factor B is the TF on the right of the network diagram. In some topologies A and B and positively correlated (left panel), whereas they are negatively correlated in other topologies (right panel). Colored dots denote the stable steady states. Colored lines connect states of their corresponding topologies. The colors of the cell states match the illustration in Figure 1. The colors of the lines denote different representative models. z-score is calculated by shifting the mean of each four attractors to 0 and then normalizing the four data points to unit variance data. B. Example phase planes for two minimum topologies (Type I and Type II respectively). In each case, four out of the seven steady states (intersections denoted by solid dots) are stable. All models shown in this figure are built with multiplicative form of Hill functions.

**Figure S5. Overlaid four attractors for each of the 559 topologies from the T cell network that produce 4-attractor systems.** Colored dots denote the stable steady states. Colored lines connect states of their corresponding topologies. The colors of the cell states match the illustration in Figure 1. The colors of the lines denote different representative models. z-score is calculated by shifting the mean of each four attractors to 0 and then normalizing the four data points to unit variance data. All models shown in this figure are built with multiplicative form of Hill functions.

**Figure S6. Comparison of three types of network topologies.** Histogram shows distributions of the numbers of topologies from the entire complexity atlas (Figure 3C) over the space of parameter sets that generate the four-attractor systems per 10^6^ random parameter sets. Distributions are separately shown for three types of motifs. Red: Type I motif. Blue: Type II motif. Green: Hybrid motif.

**Figure S7. Landscape and corresponding minimum action paths (MAPs) for the T cell developmental network in the PU.1-BCL11B state space.** White solid lines represent the MAP from ETP state to DN2a, DN2b, and DN3 states. Magenta solid lines represent the MAP from DN3 to DN2b, DN2a, and to ETP state. Dashed lines represent the direct MAP from ETP to DN3 and from DN3 to ETP states, respectively.

**Figure S8. Landscape and corresponding minimum action paths (MAPs) for the T cell developmental network in the PU.1-GATA3 state space.** White solid lines represent the MAP from ETP state to DN2a, DN2b, and DN3 states. Magenta solid lines represent the MAP from DN3 to DN2b, DN2a, and to ETP state. Dashed lines represent the direct MAP from ETP to DN3 and from DN3 to ETP states, respectively.

**Figure S9. Landscape changes as Notch signal increases.** As the Notch signal (N) increases, the landscape change from a quadristable (four stable states coexist), to tristable (DN2a, DN2b and DN3), to bistable (DN2b and DN3) and finally to a monostable DN3 state.

**Figure S10. Quasi-energy landscape for the *Bcl11b* knockout condition.** With the Bcl11b knockout (kB=0), the landscape changes from a quadristable (four stable states coexist), to a bistable (ETP and DN2a) state.

**Figure S11. Venn diagram of four types of network motifs that can produce four attractors with up to three TFs.** Red and blue areas correspond to Type I and Type II motifs shown in Figure 2B. Green area corresponds to motifs that contain both Type I and Type II networks. Orange area corresponds to motifs that can only produce four unordered attractors, in which the concentrations of the TFs are non-monotonically correlated. Numbers in the diagrams denote the total numbers of non-redundant topologies for each type. The Type II (blue) and Hybrid (green) motifs can produce both ordered and unordered 4-attractor systems, depending on the choice of parameters.

**Figure S12. Enrichment motifs containing varying numbers of positive feedback loops similar to Type I motif.** Top panel: total occurrences of various types of motifs in the T cell network. Middle panel: empirical p-values (middle panel) for these motifs with a background network population. Bottom panel: an illustration of the p-values with the distributions of the background population. Each motif has *n* (0 < *n* < 9) positive feedback loops, all of which share at least one TFs in the motif. Type I motif is a special case of such motifs withn = 3. Random networks were obtained by assigning random regulations (positive, negative or none) between each pair of TFs. 10^5^ random networks were generated. Empirical p-values were obtained by counting the number of the random networks with the motifs not less than those in the T cell network. See Methods for details of the p-value definition. Distributions of motif frequencies obtained from the random networks are shown in the bottom panel. The yellow vertical bars represent the number of occurrences in the T cell network. The right-tail areas defined by the vertical bars correspondto the shown in the middle panel (blue bars).

